# Domain-general conflict monitoring predicts neural and behavioral indices of linguistic error processing during reading comprehension

**DOI:** 10.1101/2021.01.06.425590

**Authors:** Trevor Brothers, Margarita Zeitlin, Arim Choi Perrachione, Connie Choi, Gina Kuperberg

## Abstract

The ability to detect and respond to linguistic errors is critical for successful reading comprehension, but these skills can vary considerably across readers. In the current study, healthy adults (age 18-35) read short discourse scenarios for comprehension while monitoring for the presence of semantic anomalies. Using a factor analytic approach, we examined if performance in non-linguistic conflict monitoring tasks (Stroop, AX-CPT) would predict individual differences in neural and behavioral measures of linguistic error processing. Consistent with this hypothesis, domain-general conflict monitoring predicted both readers’ end-of-trial acceptability judgments and the amplitude of a late neural response (the P600) evoked by linguistic anomalies. Interestingly, the influence on the P600 was non-linear, suggesting that online neural responses to linguistic errors are influenced by both the effectiveness and efficiency of domain-general conflict monitoring. These relationships were also highly specific and remained after controlling for variability in working memory capacity and verbal knowledge. Finally, we found that domain-general conflict monitoring also predicted individual variability in measures of reading comprehension, and that this relationship was partially mediated by behavioral measures of linguistic error detection. These findings inform our understanding of the role of domain-general executive functions in reading comprehension, with potential implications for the diagnosis and treatment of language impairments.

## Introduction

Comprehension is a goal-directed activity that allows us to infer an abstract message from strings of perceptual inputs. However, this goal is sometimes interrupted by the presence of linguistic errors. For example, in natural conversation, one in ten utterances contain some sort of speech error (Nakatani & Hirschberg, 1994), many of which are never corrected by the speaker (Levelt, 1983; Nooteboom, 1980). While error monitoring is a critical component of skilled comprehension, individuals can vary considerably in their ability to detect linguistic errors during online processing (Ehrlich, Remond, Tardieu, 1999; Garner, 1980; Hacker, 1997; Wagoner, 1983; Yudes, Macizo, Morales & Bajo, 2013). In the current study, we tested the hypothesis that linguistic error processing in healthy adult readers is affected by individual differences in domain-general *conflict monitoring*. Specifically, we asked whether the ability to monitor and resolve conflicts in non-linguistic tasks can predict *neural* and *behavioral* measures of linguistic error processing, and whether this, in turn, has consequences for comprehension success.

Language errors are common in everyday comprehension and can arise from multiple sources. For example, overt errors occur frequently in natural speech and informal written communication, including mispronunciations, typos, and word substitutions. Even in carefully edited texts, *internal* processing errors can also arise on the part of the comprehender. For example, readers regularly encounter linguistic ambiguities that result in temporary misinterpretations (Altmann, 1998), while “slips of the eye” can lead readers to incorrectly recognize one word as another (Gregg & Inhoff, 2016; Kaufman & Obler, 1995). Because of these errors, comprehenders face a signal detection problem: was the current sentence understood as intended, or did it contain an error in need of reprocessing? In order to diagnose an error, readers must detect a *conflict* between the linguistic input and their current communication model – an internal model that encodes their high-level assumptions about the communicator and the broader linguistic environment (cf. Frank & Goodman, 2012; Degen, Tessler & Goodman, 2015; see Kuperberg, Brothers & Wlotko, 2020, for recent discussion). Incoming words can conflict with a reader’s communication model at multiple levels of representation, including orthography, syntax, and semantics. For example, most readers can rapidly infer that the sentence “*He gave the candle the girl*” contains a semantic error (Gibson, Bergen & Piantadosi, 2013) because the meaning of this sentence strongly conflicts with the reader’s model of what is possible in the real world.

Once a conflict is detected, comprehenders can engage in additional compensatory behaviors like re-reading, which allow them to re-analyze the input and potentially recover a text’s intended meaning (Baker & Anderson, 1982; Carpenter & Daneman, 1981; Helder, Van Leigenhorst & van den Broek, 2016; Wagoner, 1983). For example, eye tracking studies have shown that highly implausible or syntactically unlicensed continuations are associated with longer fixation times (Kinnunen & Vauras, 1995; Traxler, Foss, Seely, Kaup & Morris, 2000; Warren & McConnell, 2007; Warren, Milburn, Patson & Dickey, 2015) and more frequent regressions to earlier parts of the text (Ni et al., 1998; Rayner, Warren, Juhazs & Liversedge, 2002; Zabrucky & Ratner, 1989). Moreover, when readers are unable to make regressive saccades, their comprehension performance suffers, particularly in sentences that require semantic or syntactic re-analysis (Schotter, Tran & Rayner, 2014; Metzner, von der Malsburg, Vasishth & Rosler, 2017). This suggests that linguistic error detection and compensatory behaviors play a causal role in supporting text understanding. Indeed, in younger readers, the ability to consciously detect and respond to language errors is considered a central component of skilled reading (Baker & Anderson, 1982; Cain, Oakhill, & Bryant, 2004; Oakhill, Hartt, & Samols, 1996; Wagoner, 1983), and these *comprehension monitoring* skills are associated with improvements in reading comprehension across development (Oakhill, & Cain, 2012).

In studies measuring event-related potentials (ERPs), linguistic anomalies have been associated with a late positive-going neural response — the *P600* – which is maximal from 600-1000ms over posterior electrode sites. While this response was initially associated with the processing of syntactic anomalies and ambiguities (Hagoort, Brown & Groothusen, 1993; Osterhout & Holcomb, 1992), later studies showed that P600s are elicited by a wide range of linguistic violations, including misspellings (Bulkes, Christianson & Tanner, 2020; Vissers, Chwilla & Kolk, 2006) and semantic anomalies (Munte, Heinze, Matzke, Wieringa & Johannes, 1998; Kuperberg; Sitnikova, Caplan & Holcomb, 2003; Szewczyk & Schriefers, 2011). For example, a robust P600 is evoked by semantically anomalous sentence continuations (e.g. *Every morning for breakfast the eggs would *<u>plant</u>*…), but not by mildly implausible continuations (e.g. *Every morning for breakfast the boys would <u>plant</u>*…) (Kuperberg, Sitnikova, Caplan & Holcomb, 2003, 2007; Paczynski & Kuperberg, 2012; van de Meerendonk, Kolk, Vissers & Chwilla, 2010). Importantly, simply detecting a semantic anomaly is not sufficient to generate this component; to produce a P600, readers must consciously perceive the input as a *comprehension error*, that is, they must be engaged in deep comprehension, and, as such, have established a communication model that conflicts with the input (Brothers, Wlotko, Warnke & Kuperberg, 2020, Kuperberg, Brothers & Wlotko, 2020)^1^. Once a conflict has been detected, readers can engage in additional second-pass mechanisms, such as *reanalysis*, in order to determine if a word was correctly perceived the first time around (see van de Meerendonk, Kolk, Chwilla & Vissers, 2009 for a discussion).^2^

The ability to monitor for conflict is thought to play a role in behavioral regulation in a wide range of non-linguistic domains (Botnivick, Braver, Barch, Carter & Cohen, 2001). In the cognitive control literature, the term *conflict monitoring* has been used to refer both to the automatic detection of conflict between competing responses (e.g. Yeung, Botvinick & Cohen, 2004), as well as the ability to consciously detect and respond to conflicts between environmental inputs and an internal model of the current task (e.g. Yu, Dayan & Cohen, 2009). These ‘conflict signals’ can then be used to regulate behavior, by inhibiting prepotent behavioral responses or selectively attending to relevant environmental inputs.^3^ For example, in the well-known Stroop task, the detection of conflict between well-learned aspects of the input (the meaning of the word “RED”) and a goal-relevant model that describes the task (identifying font colors), results in a reactive re-allocation of attention to prevent incorrect responses.

To summarize thus far, during language comprehension, the detection of a conflict between the linguistic input and a comprehension-relevant communication model is thought to trigger the P600, which may reflect re-analysis or reallocation of attention to the prior linguistic input. Analogously, in non-linguistic tasks, the detection of conflict between the input and a task-relevant internal model is thought to trigger a reallocation of attention to goal-relevant aspects of the input, in order to prevent behavioral errors. Despite these apparent similarities, it is unclear whether individual differences in domain-general conflict monitoring influence the online processing of linguistic errors in healthy adult readers.

Although there has been little systematic research into the relationship between the P600 and domain-general conflict monitoring, some previous ERP studies have examined its links with a related construct — working memory capacity, which is often measured using ‘complex span’ tasks (Daneman & Carpenter, 1980). The results of these studies are somewhat mixed. In one study (Nakano, Saron & Swaab, 2010), high-span participants showed a robust P600 response to semantically anomalous verbs while low-span participants showed an ERP effect of the opposite polarity — an N400, which has been linked to the difficulty of lexico-semantic retrieval (Kutas & Federmeier, 2011). A reading comprehension study by Oines, Miyake, and Kim (2018) found a similar pattern: verbal (but not spatial) working memory capacity predicted the same trade-off relationship between P600 and N400 responses. On the other hand, several other studies report *no* significant relationships between working memory capacity and P600 amplitudes, either in response to semantic anomalies (Ye & Zhou, 2008; Kos, van den Brink & Hagoort, 2012; Zheng & Lemhofer, 2019), or syntactic anomalies (Tanner & van Hell, 2014; Tanner, 2019).

The results of these prior studies are somewhat difficult to interpret, both due to inconsistent findings and the multi-factorial nature of working memory tasks. For example, performance in complex span tasks indexes a range of cognitive abilities, including memory storage, domain-specific verbal knowledge, and the inhibition of task irrelevant information (Bayliss, Jarrold, Gunn & Baddeley, 2003; McCabe, Roediger, McDaniel, Balota & Hambrick, 2010). Therefore, based on the previous literature, it is unclear which of these skills are most relevant for detecting linguistic errors (see Vuong & Martin, 2013 for a discussion).

### The present study

The goal of the present study was to systematically examine the relationship between domain-general conflict monitoring and neural/behavioral measures of linguistic error processing in healthy adult readers. We recorded ERPs as participants read short discourse contexts for comprehension and monitored for the presence of semantic anomalies. To assess individual differences in linguistic error processing, we examined the P600 effect evoked by semantically anomalous (versus plausible) words, as well as participants’ end-of-trial plausibility judgments (d’).

To examine the role of domain-general conflict monitoring, the same group of participants completed two conflict monitoring tasks: the *Stroop* task (MacLeod, 1991) and a fast-paced version of the *AX Continuous Performance Task* (AX-CPT; Servan-Schriber, Cohen & Steingard, 1996). These tasks were administered as part of a comprehensive neuropsychological battery, which also assessed individual differences in working memory capacity, verbal knowledge, and processing speed. We carried out a factor analysis to isolate latent ‘factor scores’ for each of these cognitive domains, and these predictor variables were used to address two main theoretical questions.

First, does monitoring for linguistic errors during comprehension rely on domain-general conflict monitoring mechanisms? If so, then individual differences in domain-general conflict monitoring should predict both neural and behavioral responses to semantic errors during reading, even after controlling for variability in working memory, verbal knowledge, and processing speed. Here, we predicted that individuals with higher domain-general conflict monitoring scores would show higher behavioral accuracy and larger P600 amplitudes in response to semantic anomalies. We also considered an alternative “efficiency hypothesis” that would produce *dissociations* between behavioral and neural measures (Gray et al., 2005; Haier, Siegel, Tang, Abel & Buchsbaum, 1992; Rypma et al., 2006). Specifically, individuals with better conflict monitoring abilities may resolve linguistic conflicts more efficiently, resulting in higher behavioral accuracy and smaller P600 responses overall.

Finally, we also examined the relationship between domain-general conflict monitoring and participants’ ability to understand and answer questions about a text (*reading comprehension*). As noted earlier, monitoring and responding to linguistic errors is thought to be an important subcomponent of reading comprehension in both adult and developing readers (Cain, Oakhill & Bryant, 2004; Garner, 1980; Wagoner, 1983). However, while it is well-established that working memory capacity and verbal abilities are important predictors of reading comprehension performance (Conway & Engle, 1996; Daneman & Carpenter, 1980; Daneman & Merikle, 1996; Cromley, Snyder-Hogan & Luciw-Dubas, 2010; Freed, Hamilton & Long, 2017; Stanovich & Cunningham, 1992), it is less clear whether domain-general conflict monitoring plays an independent role (cf. Christopher et al., 2012; McVay & Kane, 2012). In the present study, we predicted that non-linguistic conflict monitoring would account for additional variance in reading comprehension performance, and that this effect of domain-general conflict monitoring would be mediated by individual differences in linguistic error detection.

## Methods

### Participants

The current study included data from 77 participants (43 female) between the ages of 18 and 35 (mean: 23.5). This sample included 39 Tufts University undergraduates and 38 community-dwelling adults from the Boston metropolitan area, who were recruited using online advertisements. Participants were right-hand dominant (Oldfield, 1971), had no significant exposure to languages other than English before the age of five, and had no history of head injury or psychiatric/neurological diagnoses. Participants received course credit or were compensated for their participation. All protocols were approved by Tufts University Social, Behavioral, and Educational Research Institutional Review Board. All participants in this sample completed both the primary ERP study and a separate two-hour session that included a range of behavioral individual differences measures (see below).

### ERP Study

#### Experimental design and linguistic stimuli

The experimental stimuli examined here consisted of 100 three-sentence discourse scenarios. In each scenario, the first two sentences introduced a coherent discourse context (*The lifeguards received a report of sharks right near the beach. Their primary concern was to prevent any incidents in the sea…*). The third sentence contained either an animate-constraining or inanimate-constraining verb followed by a critical noun that was either plausible (*Hence, they cautioned the <u>trainees</u>*…), or semantically anomalous (*Hence, they cautioned the <u>drawer</u>*…). To create the anomalous versions of each scenario, the same animate and inanimate critical nouns were counterbalanced across contexts. Plausible and semantically anomalous nouns were matched in both lexical predictability (<1% cloze; Taylor, 1953) and mean cosine semantic similarity to the preceding context (latent semantic analysis, word-to-document similarity, *t* < 1; Landauer & Dumais, 1997). This is because both these factors are known to influence the amplitude of the N400 (Kutas & Federmeier, 2011). We then verified that the two conditions differed in plausibility by recruiting a separate group of participants from Amazon Mechanical Turk and asking them to rate each discourse scenario on a seven-point scale (7 = “makes perfect sense”, 1 = “makes no sense at all”). As expected, there were clear differences in plausibility ratings between the two conditions (plausible scenarios: M = 5.5, SD = 0.9, anomalous: M = 1.9, SD = 0.6). Additional information on the creation and norming of these stimuli are described in previous studies (Kuperberg, Brothers & Wlotko, 2020; Brothers, Wlotko, Warnke & Kuperberg, 2020).

Plausible and anomalous scenarios were counterbalanced across lists, ensuring that participants saw each discourse context and critical word only once. To maximize statistical power, data were combined from three subgroups of participants who completed slightly different versions of the main ERP experiment (for full details, see Supplementary Materials). These three versions used exactly the same presentation parameters and experimental items, and they all included equal proportions of plausible and semantically anomalous discourse scenarios. The primary differences were in the number of recording electrodes and in the relative proportions of highly constraining and non-constraining discourse scenarios. Due to differences in counterbalancing across samples, participants saw between 20 and 29 trials per condition within an experimental session.

#### Procedure

Participants sat in a sound-attenuated room, and discourse stimuli were presented on a computer monitor at a distance of 1.5 meters. The first two sentences of each scenario were presented in their entirety, one sentence at a time, which participants read at their own pace. When participants pressed a button after the second sentence, the third critical sentence appeared one word at a time in the center of the screen (450ms duration, 100ms ISI). Participants were told to read the entire discourse carefully for comprehension. Following the sentence-final word, a question mark appeared after a 1000ms delay, and participants indicated via button press whether or not the preceding discourse “made sense”. On 20% of trials, participants also answered True/False comprehension questions that probed their understanding of the whole scenario.

#### EEG preprocessing and operationalization of ERP components

EEG was recorded from a minimum of 32 scalp electrodes, arranged in a modified 10-20 system. Signals were digitized at 512 Hz with a bandpass filter of 0 Hz - 104 Hz. Offline, EEG signals were referenced to the average of the right and left mastoids, and a 0.1 - 30 Hz bandpass filter was applied. The EEG was then segmented into epochs (−200ms to 1000ms), time-locked to the onset of the critical noun. Independent component analysis was used to remove EEG artifact due to blinks, and any epochs with residual artifact were rejected (7% of trials). Artifact-free epochs were then averaged within-conditions for each participant.

Based on prior studies (Brothers et al., 2020; Kuperberg, Brothers, Wlotko, 2020), we operationalized the P600 effect as the difference in amplitude between anomalous and plausible nouns from 600-1000ms over a cluster of posterior electrode sites (Pz, P3/4, Oz, O1/2). In addition to the P600 effect, we also examined differences in the N400 component, which has been shown to vary systematically with the amplitude of the P600 effect across participants (e.g. Oines, Kim & Miyake, 2018). The N400 was operationalized as the average voltage from 300-500ms over central-parietal electrode sites (CPz, CP1/2, Pz, P3/4).

### Behavioral Assessment of domain-general cognitive functions and reading comprehension

#### Behavioral session: Task procedures

In addition to the ERP experiment, participants also completed a two-hour behavioral session on a separate day. This session included a battery of neuropsychological tasks assessing conflict monitoring, working memory, verbal knowledge, and processing speed, as well as two standardized reading comprehension assessments. Participants were tested individually in a quiet room by a trained experimenter. Nine of these tasks were administered by the experimenter, and five tasks were automated and presented on a desktop computer. The same task order was used for all participants. A short description of each task is provided below, along with scoring methods and the dependent measures of interest (for additional information, see Supplementary Materials).

#### Conflict monitoring tasks

To assess individual differences in conflict monitoring, participants completed a fast-paced version of the AX-CPT and a manual Stroop task. The AX-CPT task was first developed by Servan-Schriber, et al. (1996) to assess individual differences in proactive and reactive cognitive control. In this task, participants saw a letter cue (e.g. “A”) followed by a target (e.g. “X”). On the majority of trials (70%), participants saw a frequent cue-target pairing (”AX”), which requires a right-hand button response. On critical “AY” trials (10%), the “A” cue was followed by a different letter (e.g. “G”), and participants were required to withhold their default response and press a different key instead. The remaining 20% of trials served as control conditions, in which the target letter was preceded by a different cue (“BX” and “BY”; see Table 1).

**Table 1.** Descriptive statistics for domain-general conflict monitoring tasks

| *Variable* | *Mean* | *SD* |
| --- | --- | --- |
| *AX-CPT* |  |  |
| AX Acc. | .965 | .064 |
| AY Acc. | .807 | .177 |
| BX Acc. | .934 | .136 |
| BY Acc. | .938 | .124 |
| AX RT (ms) | 365 | 43 |
| AY RT (ms) | 476 | 50 |
| BX RT (ms) | 335 | 74 |
| BY RT (ms) | 341 | 72 |
| *Manual Stroop* |  |  |
| Congruent Acc. | .986 | .027 |
| Incongruent Acc. | .939 | .051 |
| Control Acc. | .979 | .029 |
| Congruent RT (ms) | 968 | 261 |
| Incongruent RT (ms) | 1176 | 318 |
| Control RT (ms) | 975 | 236 |

*Note.* Acc. = Accuracy, RT = Reaction Time, AX-CPT = AX Continuous Performance Task

Different versions of the AX-CPT have been developed to emphasize different aspects of cognitive control (Henderson, et al., 2012). Here we used a fast-paced version of the task with short cue-target intervals (stimulus duration: 250ms; SOA: 750ms). With this version, by de-emphasizing the role of cue maintenance, we hoped to better dissociate variability in conflict monitoring and working memory. Participants completed a short practice session with feedback, followed by 150 cue-target pairings. *The two dependent measures for this task were AY response errors and the reaction time difference between critical and control trials (AY minus AX).*

In the manual Stroop task (MacLeod, 1991), participants identified the printed color of neutral letter strings (a string of X’s), congruent color words (“blue” in blue font), and incongruent color words (“blue” in red font). There were four possible button-press responses (red, black, green, blue) and participants completed 28 trials in each condition. *The dependent measure for this task was percentage of errors in the incongruent condition.* We also calculated Stroop reaction time costs (*incongruent* minus *neutral*), but this measure was excluded from the factor analysis due to low reliability (α = .25).

#### Working memory capacity tasks

Four separate ‘complex span’ tasks were administered that assessed participants’ ability to simultaneously manipulate and store information in working memory. They included automated versions of the Reading Span and Operation Span tasks (Unsworth, Heitz, Schrock & Engle, 2005), and experimenter administered versions of the Subtract-Two Span (Salthouse, 1988) and Listening Span tasks (Daneman & Carpenter, 1980). After a short practice session, participants completed three to five trials at each span length (see Supplementary Materials). *The dependent measure in each task was the total number of items recalled in all error-free sets* (see Table 2).

**Table 2.** Mean, SD, Range, Skew, Kurtosis, and Reliability values for individual difference measures

|  |  | Mean (SD) | Range | Skew | Kurtosis | Reliability |
| --- | --- | --- | --- | --- | --- | --- |
| *Working Memory Capacity* | Operation Span | 50.0 (17.7) | 0 – 75 | -1.0 | 1.2 | .83<sup>a</sup> |
|  | Reading Span | 47.0 (18.7) | 3 – 75 | -0.5 | -0.7 | .82<sup>a</sup> |
|  | Listening Span | 29.8 (9.5) | 13 – 55 | 0.1 | -0.5 | .89<sup>a</sup> |
|  | Subtract-2 Span | 128.0 (19.4) | 70 – 169 | -0.1 | -0.1 | .92<sup>a</sup> |
| *Verbal Knowledge* | NAART | 40.4 (7.2) | 19 – 53 | -0.7 | 0.6 | .89<sup>a</sup> |
|  | Word ID | 14.0 (2.2) | 7 – 17 | -1.0 | 0.9 | .75<sup>a</sup> |
|  | Author Recognition | 19.5 (8.6) | 3 – 45 | 0.5 | 0.9 | .83<sup>b</sup> |
| *Processing Speed* | RAN (sec) | 17.0 (3.4) | 10 – 28 | 1.0 | 2.1 | .91<sup>c</sup> |
|  | RAS (sec) | 18.7 (3.5) | 11 – 30 | 0.5 | 0.7 | .91<sup>c</sup> |
|  | TOWRE Sight (sec) | 48.6 (7.4) | 72 – 35 | 0.6 | 0.4 | .91<sup>d</sup> |
| *Conflict Monitoring\** | Stroop accuracy | .934 (.051) | .79 – 1.0 | -0.8 | 0.3 | .62<sup>a</sup> |
|  | AY - AX cost (ms) | 110 (47) | 9 – 250 | 0.4 | 0.2 | .79<sup>b</sup> |
|  | AY accuracy | .807 (.177) | .40 – 1.0 | -0.8 | -0.3 | .83<sup>a</sup> |

a: Cronbach's alpha, b: Spearman-Brown split half, c: Howe, et al., 2006. d: Torgesen, Wagner, &
Rashotte, 2012, \*Stroop RT costs were excluded from the factor analysis due to low reliability ( $\alpha = 0.25$ )

#### Verbal knowledge tasks

Participants completed three tasks assessing their reading experience and verbal knowledge (Uttl, 2002; Woodcock & Johnson, 1989; Acheson, Wells & MacDonald, 2008). In the North American Adult Reading Test (NAART) and Woodcock-Johnson Word Identification task (WORD ID), participants read lists of low frequency words aloud (*quadruped*, *leviathan*). *In both tasks, the dependent measure was the number of correct pronunciations*, which was scored by two independent raters using audio recordings. In the Author Recognition Task (ART), participants selected known authors from a list of 65 famous authors and 65 non-famous foils. *The dependent measure was the number of correct identifications minus false alarms.*

#### Processing speed tasks

To measure individual differences in verbal processing speed, three rapid naming tasks were administered: Rapid Automatized Naming (RAN), Rapid Alternating Stimulus (RAS), and the Test of Word Reading Efficiency (TOWRE) Sight Word Efficiency task (Denkla & Rudel, 1976; Wolf, 1986; Torgesen, Wagner, & Rashotte, 2012). In these tasks, participants read printed lists of letters, letters and digits, or high-frequency words as quickly as possible without making mistakes. Because these tasks only provided a single outcome measure, measures of test-retest reliability were obtained from prior studies (Howe, et al., 2006; Torgesen, Wagner, & Rashotte, 2012). *The dependent measure for these tasks was time to completion, measured to the nearest second.*

#### Reading comprehension assessments

In addition to these primary neuropsychological measures, participants also completed two reading comprehension tasks: a) the comprehension portion of the Kauffman Test of Educational Achievement (KTEA), which involves reading passages and answering multiple choice comprehension questions, and b) the Woodcock Reading Mastery Test (WRMT), which involves reading short passages and filling in a missing word (Singer, Lichtenberger, Kaufman, Kaufman & Kaufman, 2012; Woodcock, 1973). *The dependent measure for these assessments was the total number of correct responses.*

### Preprocessing and Factor analysis

The neuropsychological battery described above generated 13 performance measures in total. Prior to analysis, any scores more than three standard deviations above the group mean were replaced with this cutoff value (1.2% of scores). Average performance, excess skew and kurtosis, and the reliability for each dependent measure are shown in Table 2. As expected, these tasks showed acceptable to excellent levels of reliability (α = .62 – .92). A correlation matrix describing the relationships among these measures is presented in Table 3.

**Table 3.** Correlation Matrix for individual differences measures (N = 77)

|  | 1 | 2 | 3 | 4 | 5 | 6 | 7 | 8 | 9 | 10 | 11 | 12 |
| --- | --- | --- | --- | --- | --- | --- | --- | --- | --- | --- | --- | --- |
| 1) O-SPAN | - |  |  |  |  |  |  |  |  |  |  |  |
| 2) R-SPAN | **.65** | - |  |  |  |  |  |  |  |  |  |  |
| 3) L-SPAN | **.49** | **.49** | - |  |  |  |  |  |  |  |  |  |
| 4) SUBT-2 | **.52** | **.54** | **.62** | - |  |  |  |  |  |  |  |  |
| 5) ART | .12 | **.33** | .22 | .16 | - |  |  |  |  |  |  |  |
| 6) Word_ID | **.41** | **.43** | **.54** | **.50** | **.44** | - |  |  |  |  |  |  |
| 7) NAART | **.42** | **.50** | **.42** | **.52** | **.62** | **.78** | - |  |  |  |  |  |
| 8) RAS | **.30** | **.27** | .21 | **.27** | .00 | **.24** | **.25** | - |  |  |  |  |
| 9) RAN | **.34** | **.26** | **.29** | **.45** | .08 | **.38** | **.35** | **.76** | - |  |  |  |
| 10) TOWRE | .21 | **.32** | **.28** | **.26** | .00 | **.27** | **.25** | **.54** | **.48** | - |  |  |
| 11) Stroop<sub>acc</sub> | .17 | .22 | .18 | **.26** | -.02 | **.26** | **.24** | -.09 | -.01 | .10 | - |  |
| 12) AY<sub>acc</sub> | .14 | .20 | **.29** | .14 | .15 | **.46** | **.32** | .17 | .19 | **.24** | **.24** | - |
| 13) AY RT<sub>cost</sub> | .05 | .08 | .12 | .10 | .05 | **.29** | .17 | -.12 | .06 | .11 | **.23** | **.56** |

Reaction times measures (RAS, RAN, TOWRE, AY RT<sub>cost</sub>) were reverse coded, with positive values indicating better performance. All bolded correlations ( $r > .22$ ) were significant at $p < .05$ .

Using these scores, we conducted an exploratory factor analysis, which aimed to estimate the underlying latent constructs corresponding to our four cognitive abilities of interest. This analysis was carried out using SPSS 25. In order to maximize the independent variance of each latent variable, the factor analysis was fit using a Maximum Likelihood with a Varimax rotation. The total number of factors was selected using parallel analysis (Horn, 1965), and factor scores were calculated for each participant using the Bartlett regression method (Grice, 2001).

Consistent with the hypothesized structure of our neuropsychological battery, this exploratory factor analysis yielded a four-factor solution (see Table 4). Factor 1 explained 19% of the total variance and loaded strongly on measures of *working memory capacity*. Factor 2 loaded on measures of *processing speed* (16%), Factor 3 loaded on measures of *verbal knowledge* (13%), and Factor 4 loaded on measures of *conflict monitoring* (11%). Participant-specific factor scores were then used as predictor variables in a series of multiple regression analyses designed to test our *a priori* hypotheses, described below.

**Table 4.**
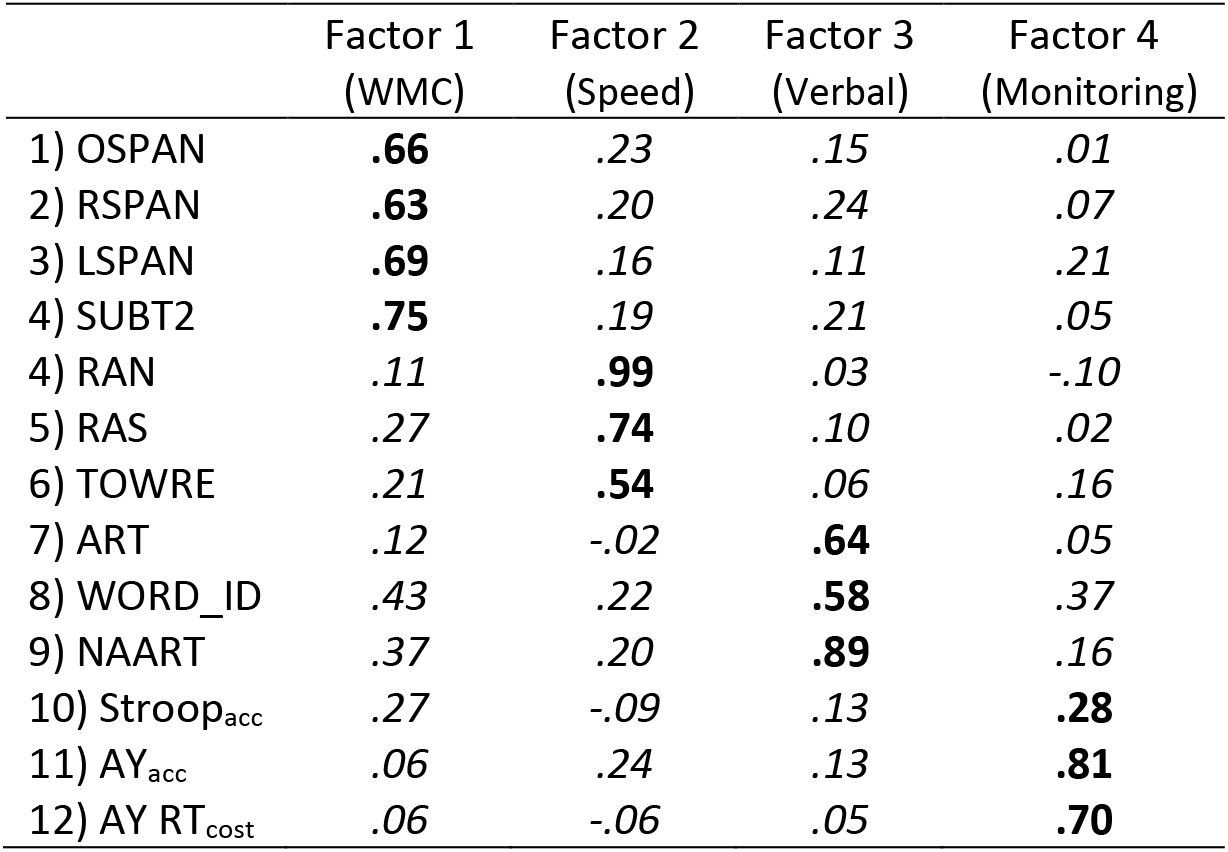
Pattern matrix for the four-factor solution obtained in an exploratory factor analysis

### Hypothesis testing

To test our primary hypotheses, we examined individual differences in two measures of linguistic error processing: (1) *behavioral detection of semantic errors*, as indexed by participants’ acceptability judgments at the end of each scenario in the ERP experiment — d’ scores; and (2) *neural semantic error processing*, as indexed by the magnitude of the P600 effect to anomalous vs. plausible critical words. For each of these measures, we first examined the influence of domain-general conflict monitoring using simple regression. We then carried out a multiple regression analysis, which included the conflict monitoring factor score and three other performance scores derived from our factor analysis. These multiple regression analyses enabled to us to determine whether working memory, verbal knowledge, or processing speed predicted additional variability in our dependent measures, and whether any significant effects of conflict monitoring remained after controlling for these other factors. In these regression analyses, we included both linear and quadratic effects of each predictor in order to capture potential non-linear contributions of each cognitive construct. We included these predictors because some prior studies have shown non-linear relationships between executive functions and the magnitude of evoked neural responses (see Luna, Padmanabhan & O’Hearn, 2010, Yarkoni & Braver, 2010 for reviews).

Finally, we performed a multiple regression analysis to ask whether domain-general conflict monitoring predicted variance in reading comprehension ability, beyond the effects of working memory and verbal knowledge. An additional mediation analysis tested whether any relationship between conflict monitoring and reading comprehension could be partially explained by individual differences in behavioral measures of linguistic error detection. In all multiple regression analyses, we standardized all predictors and dependent measures. This enabled us to compare the magnitude of effect sizes across analyses (*small*: b = 0.1; *medium*: b = 0.3; *large*: b = 0.5; Cohen, 1992; see Supplementary Materials).

## Results

### ERP Study

#### Behavioral detection of semantic errors: acceptability judgments

In the main ERP experiment, participants were able to categorize discourse scenarios as plausible (mean accuracy = 87%, SD = 11%) and anomalous (mean accuracy = 90%, SD = 10%). Although all participants performed above chance (d’ = 2.6), there was also considerable variability in behavioral sensitivity across participants (SD = 0.7, range: 1.0 to 4.7, α = 0.69).

#### Neural index of semantic error processing

Relative to plausible continuations, semantically anomalous words elicited a biphasic ERP response with larger (more negative) N400 amplitudes from 300-500ms after word onset, and larger P600 amplitudes from 600-1000ms. As reported in previous studies (Kuperberg, Brothers & Wlotko, 2020) these effects were both highly significant (*N400 effect*: M = 1.1μV, SD = 2.5, t(76) = 3.96, *p* < .001; *P600 effect*: M = 2.2μV, SD = 3.3, t(76) = 5.94, *p* < .001) and were maximal at posterior electrode sites.

To determine the split-half reliability of this P600 anomaly effect, we calculated the amplitude of this ERP difference (*anomalous* vs. *plausible*) for each participant, separately for even and odd trials. After applying the Spearman-Brown prophecy formula, the P600 effect was moderately reliable across participants (ρ = 0.61).

### Multiple regression analyses

#### A linear relationship between domain-general conflict monitoring and behavioral measure of semantic error detection

Consistent with our main hypothesis, we observed a significant linear relationship between domain-general conflict monitoring and participants’ ability to detect semantic anomalies, as indexed by their behavioral responses at the end of each scenario (b = 0.29, t = 2.67, *p* = 0.009; see Figure 1). This linear effect of conflict monitoring remained significant in a multiple regression analysis that included all four factor scores (b = 0.27, t = 2.02, *p* = 0.047). Participants with higher conflict monitoring abilities showed greater behavioral sensitivity to semantic errors, while working memory, verbal knowledge, and processing speed contributed no additional unique variance (see Table 5). We also replicated this effect of monitoring effect in an independent group of participants, using a different set of linguistic stimuli (see Supplementary Materials).

**Figure 1.**
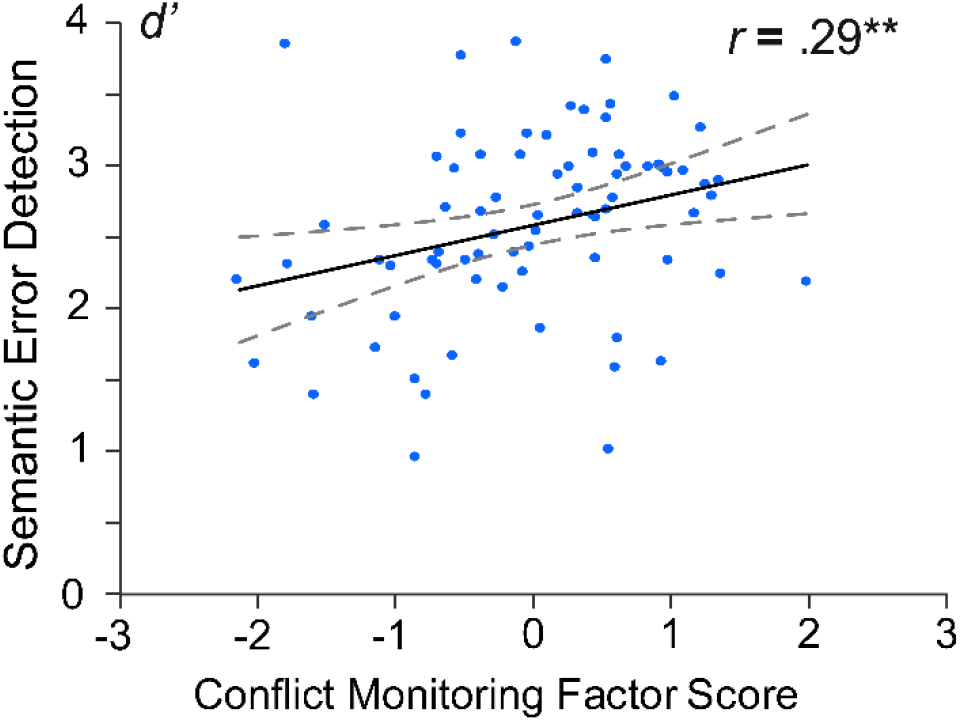
The linear relationship between domain-general conflict monitoring and semantic error detection during comprehension. Dotted lines represent 95% confidence intervals. ** *p* < .01

**Table 5.**
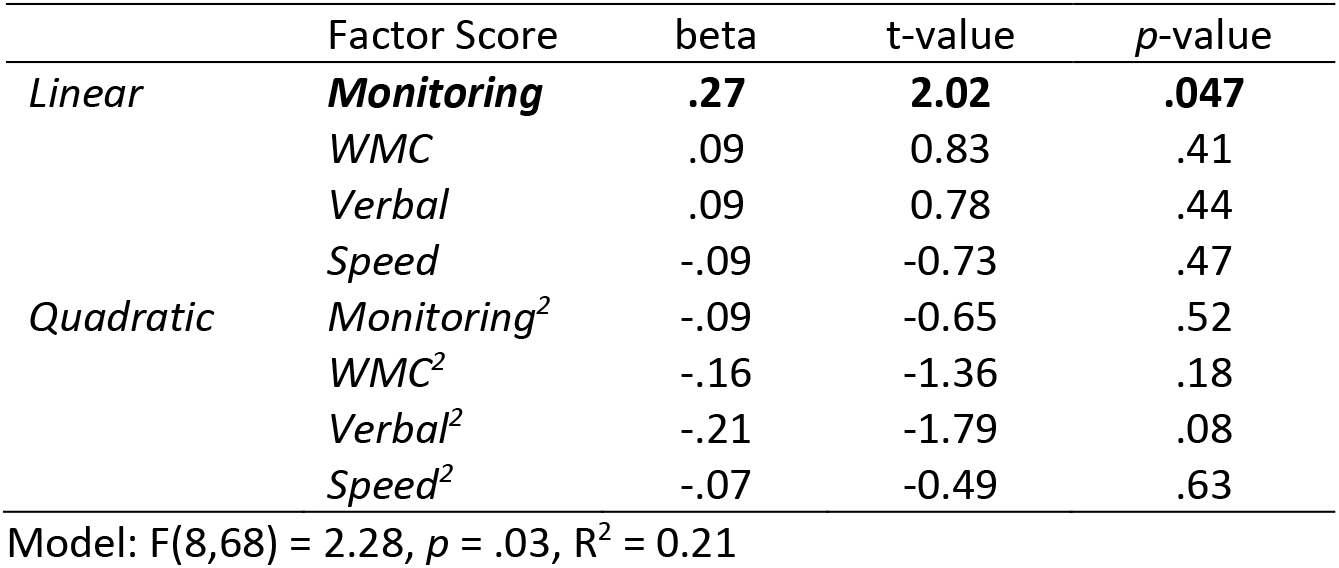
Multiple regression analysis predicting semantic error detection (d’)

|  | Factor Score | beta | t-value | p-value |
| --- | --- | --- | --- | --- |
| *Linear* | **Monitoring** | **.27** | **2.02** | **.047** |
|  | *WMC* | .09 | 0.83 | .41 |
|  | *Verbal* | .09 | 0.78 | .44 |
|  | *Speed* | -.09 | -0.73 | .47 |
| *Quadratic* | *Monitoring*<sup>2</sup> | -.09 | -0.65 | .52 |
|  | *WMC*<sup>2</sup> | -.16 | -1.36 | .18 |
|  | *Verbal*<sup>2</sup> | -.21 | -1.79 | .08 |
|  | *Speed*<sup>2</sup> | -.07 | -0.49 | .63 |

Model: $F(8,68) = 2.28$ , $p = .03$ , $R^2 = 0.21$

#### An inverse U-shaped relationship between domain-general conflict monitoring and the P600 effect

We next examined the effect of domain-general conflict monitoring on the magnitude of the P600 effect evoked by semantically anomalous (versus plausible) critical words. We observed no significant *linear* effect, but instead saw a robust *quadratic* relationship between these two measures (b = −0.40, t = −3.76, *p* < .001, for additional evidence see Supplementary Materials). This quadratic relationship was due to relatively small P600 effects in individuals with low conflict monitoring abilities, a larger P600 effect in individuals in the middle of the scale, and a small P600 effect in participants with the strongest conflict monitoring abilities (see Figure 1). In a multiple regression analysis that included all four factors scores, the quadratic effect of domain-general conflict monitoring remained significant (b = −.34, t = −2.52, *p* = .014). Again, we saw no independent effects of working memory, verbal knowledge, or processing speed on the magnitude of the P600 effect (model R^2^ = 0.19, see Table 6).

**Table 6.** Multiple regression analysis predicting P600 effect magnitudes

|  | Factor Score | beta | t-value | p-value |
| --- | --- | --- | --- | --- |
| *Linear* | *Monitoring* | .07 | 0.51 | .61 |
|  | *WMC* | -.12 | -1.11 | .27 |
|  | *Verbal* | .01 | 0.07 | .94 |
|  | *Speed* | .14 | 1.08 | .29 |
| *Quadratic* | ***Monitoring<sup>2</sup>*** | **-.34** | **-2.52** | **.014** |
|  | *WMC<sup>2</sup>* | -.05 | -0.44 | .66 |
|  | *Verbal<sup>2</sup>* | .03 | 0.23 | .82 |
|  | *Speed<sup>2</sup>* | -.02 | -0.15 | .88 |

Model: $F(8,68) = 2.04, p = .055, R^2 = 0.19$

**Table 7.**
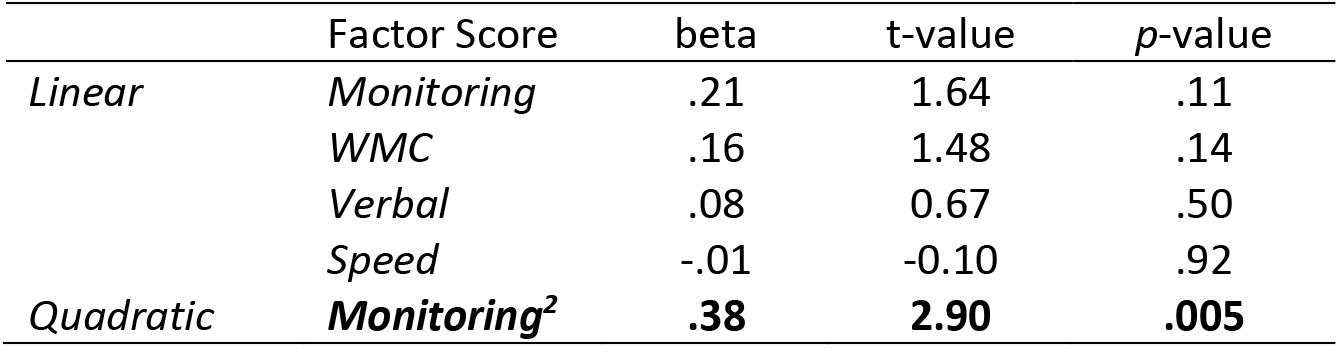
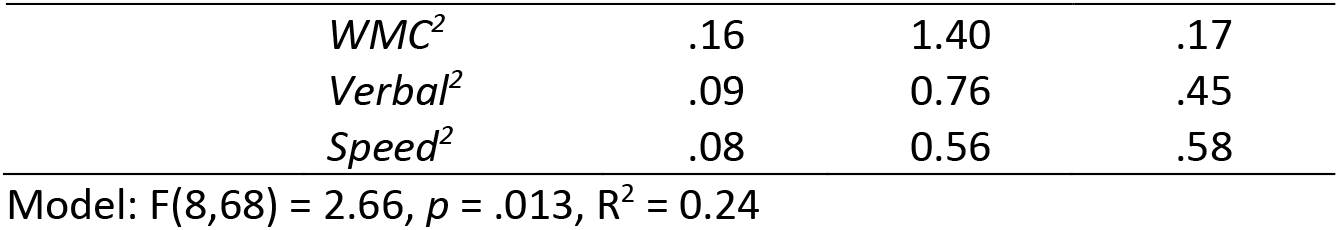
Multiple regression analysis predicting the magnitude of the N400 effect

To confirm that this quadratic effect reflected an inverse U-shaped relationship, we used an *interrupted regression* analysis to test for the presence of a change in the sign of the regression coefficient (the *two-line test*, Simonsohn, 2018). Consistent with a U-shaped function, this analysis showed a positive relationship between domain-general conflict monitoring and the P600 effect over the lower half of the range (b = 3.5, z = 3.46, *p* < .001) and a negative relationship over the upper half of the range (b = −2.4, z = −2.74, *p* = .006).

To help visualize the influence of domain-general conflict monitoring on the magnitude of the P600 effect, we split participants into three groups (High, Medium, Low) based on their conflict monitoring scores. As shown in Figure 3, individuals with intermediate scores showed the most robust P600 effects (4.0μV), while smaller P600 effects were observed in both low conflict-monitoring (1.0 μV) and high conflict-monitoring participants (1.7μV). As can be seen in Figure 3, ERP responses to plausible words were relatively constant across the three groups, and differences in the magnitude of the P600 effect were primarily driven by differential P600 responses to semantically anomalous words.

**Figure 2.**
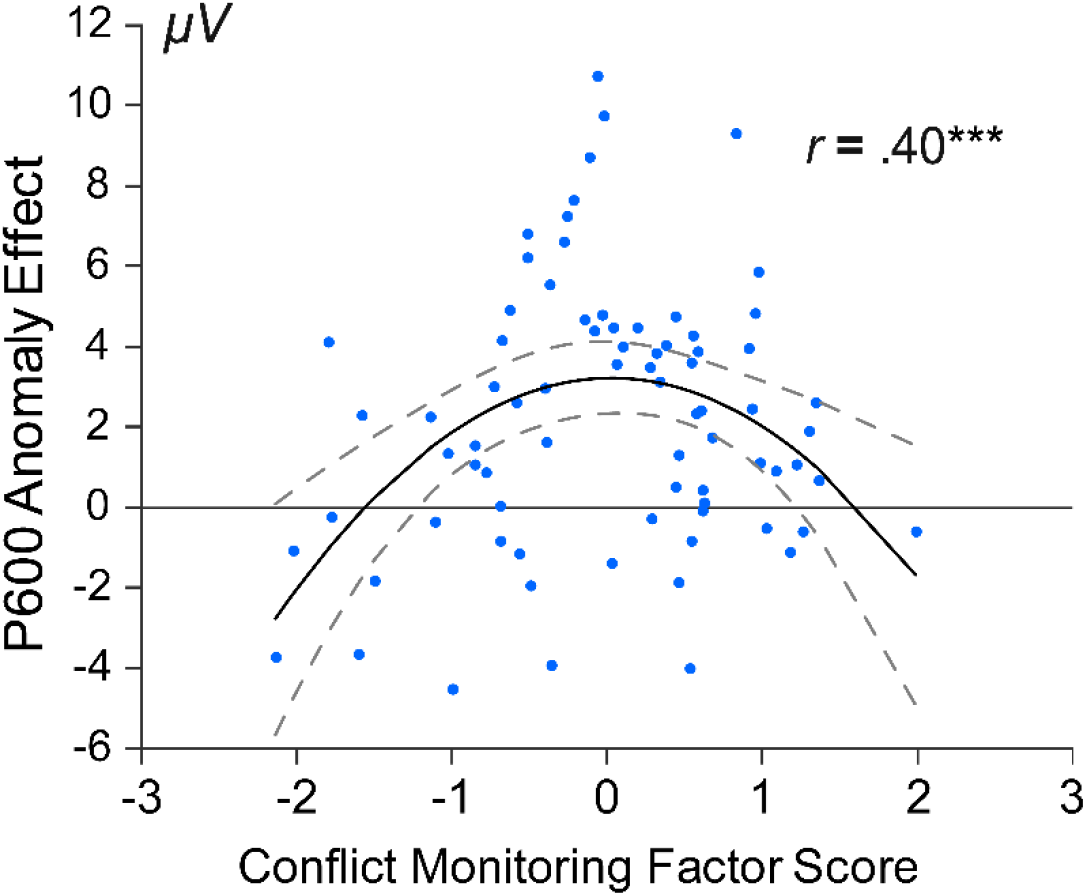
The quadratic relationship between domain-general conflict monitoring and the magnitude of the P600 effect to semantic anomalies vs. plausible control words (600-1000ms). Dotted lines represent 95% confidence intervals. *** *p* < .001

**Figure 3.**
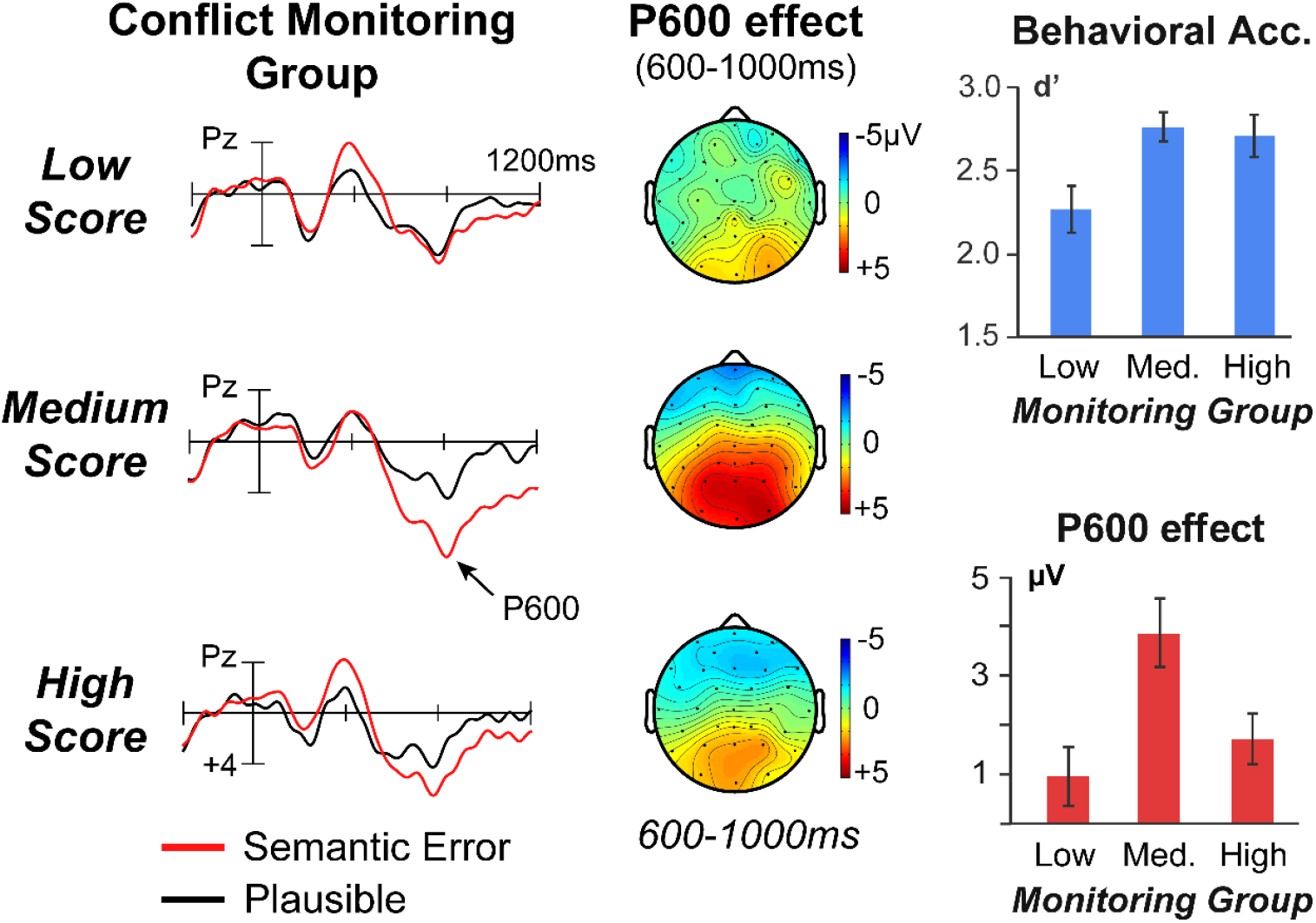
*Left*: Evoked event-related potential (ERP) responses to plausible (black) and semantically anomalous (red) critical words, plotted separately for participants with Low, Medium, and High domain-general conflict monitoring abilities. *Middle*: Topographic plots showing the magnitude and distribution of the P600 effect in each group. *Right*: Mean behavioral sensitivity to semantic anomalies (top) and mean magnitude of the P600 effect (bottom) in each of the three groups. Error bars represent ±1 SEM.

In addition, we also observed an apparent trade-off between the magnitude of the N400 and P600 effect across individuals. Consistent with prior studies (Nakano, Saron & Swaab, 2010; Oine, Myake & Kim; 2018, Tanner & van Hell, 2014; Tanner, 2019), participants with larger P600 anomaly effects showed smaller N400 differences in the 300-500ms time window (see Figure 3). Across all participants there was a significant negative correlation between the magnitude of the N400 and P600 effects to semantic anomalies, r(75) = −.60, *p* < .001.

To explore this relationship further, we conducted an additional multiple regression analysis, examining individual differences in the magnitude of the N400 effect (300-500ms). As expected, this analysis also revealed a significant quadratic effect of conflict monitoring, which was opposite in sign to that observed in the P600 time-window (b = 0.38, t = 2.90, *p* = .005). Individuals with intermediate levels of conflict monitoring showed the smallest N400 differences (−0.4 μV), and larger N400 effects were observed in both low-monitoring (1.3μV) and high-monitoring participants (2.3μV). As we note in the Discussion section, we attribute this reciprocal relationship to spatiotemporal overlap between the N400 and P600 components at the scalp surface.

#### Domain-general conflict monitoring and behavioral semantic error detection predict reading comprehension

In our final set of analyses, we examined the effects of domain-general conflict monitoring on individual differences in reading comprehension. As expected, the two measures of reading comprehension were highly correlated (r(75) = 0.57, *p* < .001), and so we combined these scores into a single index.

A multiple-regression analysis revealed that both working memory (b = 0.41, t = 4.75, *p* < .001) and verbal knowledge (b = 0.46, t = 5.26, *p* < .001) were strongly associated with reading comprehension ability (R^2^ = 0.47), replicating prior findings (Conway & Engle, 1996; Daneman & Merikle, 1996; Freed, Hamilton & Long, 2017). Importantly, we also observed an independent effect of domain-general conflict monitoring, with stronger conflict monitoring abilities predicting better reading comprehension (b = 0.20, t = 2.33, *p* = .02).

Finally, we asked whether this effect of domain-general conflict monitoring on reading comprehension could be partially explained by individual differences in linguistic error detection (d’). As shown in Figure 4, the indirect path (Conflict Monitoring → Semantic Error Detection → Comprehension) was significant (Sobel’s test = 2.06, *p* = .04), and the direct effect of conflict monitoring was partially attenuated after accounting for variability in linguistic error detection (Control: b = .14, t = 1.62, *p* = .11). This suggests that the link between domain-general conflict monitoring and comprehension relies, to some extent, on differences in the ability to detect linguistic errors.

**Figure 4.**
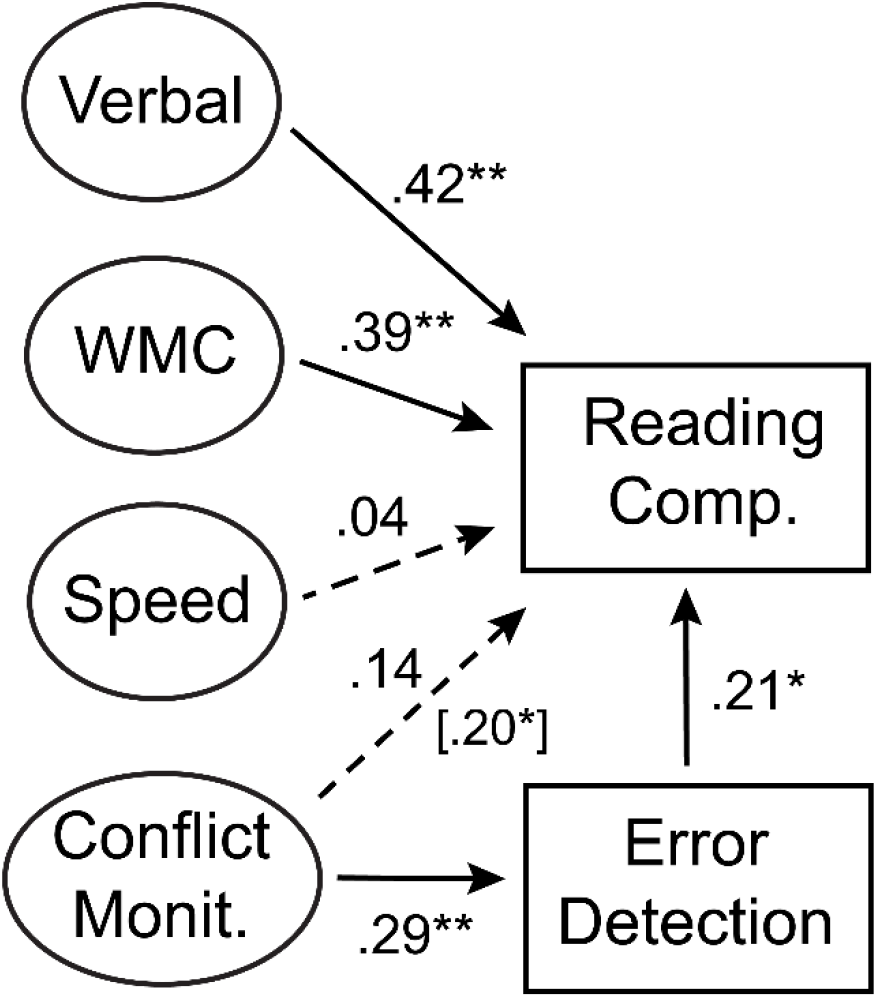
A path diagram representing the effects of verbal knowledge, working memory capacity (WMC), processing speed, and domain-general conflict monitoring on reading comprehension. The direct effect of domain-general conflict monitoring was partially attenuated after accounting for variability in semantic error detection (see text for explanation). * *p* < .05, ** *p* < .01

## Discussion

We carried out a large individual differences study to examine the role of domain-general conflict monitoring in linguistic error processing. Participants read short discourse scenarios that sometimes contained semantic anomalies, and we examined variability across readers in both conscious error detection and the amplitude of the P600 response. Participants also completed a battery of tasks assessing domain-general conflict monitoring, working memory capacity, verbal knowledge, processing speed, and reading comprehension. We found that domain-general conflict monitoring was associated with behavioral error detection (with a linear relationship) and the magnitude of the P600 effect (with an inverse U-shaped relationship). We further showed that domain-general conflict monitoring predicted variability in reading comprehension ability – above and beyond the effects of working memory and verbal knowledge – and that this relationship could be partially explained by individual differences in semantic error detection. We discuss each of these findings in more detail below. We then discuss some open questions raised by our research as well as the theoretical and practical implications of our findings.

### Domain-general conflict monitoring predicts both behavioral and neural indices of semantic error processing

The observed relationships between domain-general conflict monitoring and linguistic error processing were highly specific; that is, they could not be explained by variance in other cognitive abilities such as working memory capacity and verbal knowledge. We take this as evidence that *linguistic* error processing and *non-linguistic* conflict monitoring engage an overlapping set of cognitive processes, which involve detecting and responding to conflicts between environmental inputs and an internal mental model of the current task.

Obviously, the nature of these goal-relevant internal models differed during non-linguistic conflict monitoring and monitoring for linguistic errors during comprehension. For example, in the AX-CPT, participants were asked to respond whenever an “A” was followed by an “X”. To carry out this task efficiently and accurately, they likely engaged an internal model that represented their knowledge about the predictive relationships between cues and targets (Cohen, Barch, Carter & Servan-Schreiber, 1999; Yu, Dayan & Cohen, 2009). In contrast, during language processing, the goal was to *comprehend* the linguistic input (infer the communicator’s intended message). To achieve this goal, participants likely engaged an internal default “communication model” that represented their default linguistic and real-world knowledge (Kuperberg & Jaeger, 2016; Kuperberg, Brothers, & Wlotko, 2020). Critically, however, in both situations, participants sometimes encountered a bottom-up input that *conflicted* with this goal-relevant model (in the AX-CPT, an AY trial; in language comprehension, a semantic anomaly). We suggest that in both cases, the detection of this conflict led to a *disruption* of the default mode of processing, and triggered a reactive reallocation of attentional resources that enabled participants to achieve their goal. In the AX-CPT, this involved reallocating attention to the unexpected target letter, allowing for a shift to an alternative behavioral response (to avoid an error in action); in language comprehension, it involved reallocating attention to the critical word and prior context (*reanalysis*) in an attempt to re-establish coherence (thereby avoiding a potential error in comprehension).

Although domain-general conflict monitoring predicted both behavioral and neural measures of semantic error processing, the nature of these relationships differed: We observed a *linear* relationship with participants’ behavioral sensitivity to semantic errors (d’), and a *non-linear* (inverse U-shaped) relationship with neural measures of error processing. Specifically, the P600 effect was largest in readers with moderate domain-general conflict monitoring ability and smallest in individuals with low or high conflict monitoring ability (see Figure 3). Importantly, the same pattern of linear and non-linear effects were also observed in a separate group of participants, using a different set of linguistic stimuli (N=37, see Supplementary Materials).

Although this type of dissociation between neural and behavioral measures is relatively novel in the ERP literature, similar dissociations have previously been reported in functional magnetic resonance imaging (fMRI) studies examining different aspects of executive function (see Manoach, 2003; Luna et al., 2010, Yarkoni & Braver, 2010 for reviews). For example, in developmental studies of the *anti-saccade* task, hemodynamic responses within frontal control regions were largest in participants with intermediate behavioral performance (adolescents, ages 14-17), and smallest when anti-saccade performance was either very poor (children, age 8-13) or close to ceiling (adults, age 18-30) (Luna et al., 2001; Luna et al., 2010). Similar U-shaped response profiles have also been observed in studies that varied the difficulty of working memory demands. For example, in fMRI studies using the N-back task, large responses in control regions were observed at intermediate levels of cognitive load, while reduced activity was observed when cognitive load was minimal or when load was very high and behavioral performance began to break down (Callicott et al., 1999; Ciesielski, Lesnick, Savoy, Grant & Ahlfors, 2006; Mattay, et al, 2006; see also Vogel & Machizawa, 2004).

In these previous neuroimaging studies, inverse U-shaped responses have been interpreted as reflecting a trade-off between the *effectiveness and efficiency* of executive control mechanisms (Gray et al., 2005; Haier, Siegel, Tang, Abel & Buchsbaum, 1992; Larson, Haier, LaCasse & Hazen, 1995; Rypma et al., 2006). For example, when participants are unable to carry out a task correctly, they may fail to engage control mechanisms at all. At moderate levels of task difficulty, accurate task performance engages maximum executive resources, resulting in the largest neural activity. Finally, when participants are highly skilled in a task, they are able to carry it out with increased efficiency, resulting in reduced neural responses.

In the present study, we suggest that participants with the worst domain-general conflict monitoring performance had the most difficulty detecting semantic errors, as evidenced by their poor behavioral performance and relatively small P600 effects (see Sanford, Leuthold,Bohan & Sanford, 2010; Batterink & Neville, 2013, for evidence that only detected anomalies elicit P600 responses). In contrast, participants with intermediate domain-general conflict monitoring were better able to detect the presence of semantic anomalies and may have been more likely to engage neural resources to reanalyze the linguistic input, resulting in a robust P600 response. Finally, for participants with high conflict monitoring abilities, we believe they exhibited high accuracy and small P600 effects because their neural processing was maximally efficient. For example, these high-performing participants may have been able to efficiently categorize semantic errors as anomalous without engaging in extensive re-processing of the critical word or the prior context. Alternatively, they may needed fewer neural resources to reanalyze the input, compared to other readers.

Note, the account outlined above assumes that our behavioral measure (d’) primarily reflected comprehenders’ abilities to consciously *detect* the presence of linguistic errors, while the magnitude of the P600 effect was sensitive, not just to error detection (Sanford, Leuthold,Bohan & Sanford, 2010; Batterink & Neville, 2013) but also additional processing stages, including a reanalysis of the anomalous input and the prior context (see Brothers, Wlotko, Warnke & Kuperberg, 2020; Metzner, von der Malsburg, Vasishth & Rosler, 2017).

### N400-P600 tradeoffs

In addition to predicting the magnitude of the P600 anomaly effect, we also observed an effect of domain-general conflict monitoring on the magnitude of the N400 effect, which is thought to reflect the difficulty of lexico-semantic access/retrieval (Kutas & Federmeier, 2011). This N400 conflict monitoring effect was exactly opposite to the relationship observed for the P600, with smaller N400 differences in readers with moderate conflict monitoring ability, and larger N400 differences for individuals with either low or high conflict monitoring ability. Reciprocal relationships between N400 and P600 amplitudes have been observed in previous ERP studies examining semantic or syntactic anomalies (Nakano, Saron & Swaab, 2010; Oines, Miyake & Kim, 2018; Kos, van den Brink & Hagoort, 2012; Tanner & van Hell, 2014; Tanner, 2019). It has been suggested by some researchers that this reciprocal relationship reflects a trade-off in processing strategies as readers attempt to resolve competing linguistic constraints (Oines, Miyake & Kim, 2018). According to this trade-off account, “P600 dominant” individuals are more likely resolve semantic anomalies through structural reanalysis (*The meals were devouring −> The meals were devoured*), while “N400 dominant” individuals are more likely to attempt to retrieve the semantic features of anomalous critical words (Oines, Miyake & Kim, 2018; Kim & Osterhout, 2005).

Given that the semantic errors in the present experiment had no plausible syntactic edit (*Hence, they cautioned the drawer*… → ??), we believe that an alternative explanation is more likely. Specifically, we suggest that the negative correlation between N400 and P600 amplitudes simply reflects spatiotemporal overlap of ERP components with opposite polarities (for discussion, see Kuperberg, Kreher, Sitnikova, Caplan & Holcomb, 2007; Tanner, Goldshtein & Weissman, 2018; Brouwer & Crocker, 2017). EEG responses always reflect a combination of multiple neural generators, and ERPs with opposite polarities are known to ‘cancel out’ at the scalp surface (Luck, 2014). Because N400 and P600 responses often have similar scalp distributions and overlapping time courses, any increase in the amplitude of the P600 will result in a decrease the magnitude of the N400 (and vice-versa). We suggest that *all three* monitoring groups had greater difficulty retrieving the meaning of semantically anomalous words, resulting in an N400 effect. However, differences in conflict monitoring affected their ability to consciously detect and re-analyze these anomalies. This resulted in greater P600 responses for some participants, leading to greater amounts of component cancellation in the N400 time-window. Therefore, rather than reflecting a *cognitive* trade-off between semantic and structural processing, we believe this negative correlation simply reflected the cancellation of two independent neural generators.

### No effect of working memory capacity or verbal knowledge on semantic error processing

Unlike domain-general conflict monitoring, neither working memory capacity nor verbal knowledge were predictive of individual differences in linguistic error processing. Both these predictors showed high levels of internal validity and were strongly associated with individual differences in reading comprehension (see below). Therefore, it is unlikely that these null results reflected issues of measurement error or construct validity. Instead, they suggest that individual variability in working memory and verbal knowledge do not contribute substantially to linguistic error processing in skilled adult readers.

The absence of a relationship between the P600 effect and working memory capacity is consistent with some but not all previous findings in the literature. As noted in the Introduction, some studies have reported positive correlations between working memory and the magnitude of the P600 (Nakano, Saron & Swaab, 2010; Oines, Mikaye & Kim, 2018), while others have found no significant relationship (Kos, van den Brink & Hagoort, 2012; Zheng & Lemhofer, 2019; Tanner, 2019). In the present study, despite the inclusion of multiple span measures and a relatively large sample, we observed no significant correlations, either at the level of individual tasks (Subtract-2: *r* = .01, LSPAN: *r* = 0.06, RSPAN: *r* = −0.02, OSPAN: *r* = 0.03), or for our combined working memory factor score, (r(75) = −0.06, *p* = .58). One possibility is that this null effect can be explained by differences in the participants or linguistic stimuli across studies (e.g. discourse scenarios vs. single sentences). Another possibility is that the relationship between working memory capacity and the P600 is either non-existent, or so small that it is difficult to detect reliably in a single experiment. Ultimately, a combination of pre-registered ERP studies and a systematic meta-analysis of the literature may be needed to definitively resolve this issue.

At face value, the null relationship between verbal knowledge and linguistic error processing appears to contradict some prior studies examining second-language (L2) learners. In these studies, low-proficiency L2 learners often show absent or reduced P600 effects compared to native speakers (Osterhout, McLaughlin, Pitkänen, Frenck-Mestre & Molinaro, 2006; Zheng & Lemhofer, 2019). Moreover, the magnitude of the P600 in these groups correlates with both error detection rates and measures of L2 proficiency (Tanner, McLaughlin, Herschensohn & Osterhout, 2013; Zheng & Lemhofer, 2019). Critically, however, unlike most native English speakers, lower-proficiency L2 learners are likely to lack some of the core semantic and syntactic knowledge necessary to detect linguistic anomalies.^4^ In light of these results, our current findings suggest that, as readers reach native-like proficiency, subtle differences in verbal knowledge become less important. Instead, in this sample of native speakers, most of the variability in error monitoring performance depended on differences in non-linguistic conflict monitoring, which may regulate the successful *application* of stored linguistic knowledge during real-time comprehension.

### Domain-general conflict monitoring predicts measures of reading comprehension

In addition to examining its relationship with linguistic error processing, we were also interested in whether domain-general conflict monitoring predicted individual differences in comprehension ability, as indexed by standardized measures of reading comprehension. From previous studies, it is clear that comprehension abilities in adult readers vary as a function of working memory capacity (Conway & Engle, 1996; Daneman & Carpenter, 1980; Daneman & Merikle, 1996) and verbal knowledge (Cromley, Snyder-Hogan & Luciw-Dubas, 2010; Freed, Hamilton & Long, 2017; Stanovich & Cunningham, 1992). In the current study, we replicated these findings: working memory capacity and verbal knowledge uniquely accounted for 16% and 20% of the variance in reading comprehension performance.

Importantly, even after controlling for differences in working memory, verbal knowledge, and processing speed, we found that approximately 4% of the additional variance in reading comprehension ability could be explained by individual differences in domain-general conflict monitoring. Additional analyses suggested that this conflict monitoring effect was partially mediated by participants’ ability to detect linguistic errors. To explain these relationships, we hypothesize that skilled readers continually monitor for potential errors in comprehension in order to maintain coherence (van de Meerendonk, Kolk, Chwilla & Vissers, 2009). If a reader fails to detect these conflicts, or is unable to resolve them efficiently, they may continue with an incorrect or internally contradictory interpretation of a text’s meaning (Oakhill, Hartt & Samols, 1996), which may have negative, down-stream consequences for comprehension. On this account, *individuals with poor domain-general conflict monitoring abilities are poorer comprehenders because they are less able to detect and resolve to their own processing errors.*

Notably, domain-general conflict monitoring accounted for a smaller proportion of variance in comprehension performance (4%) compared to the effects of working memory capacity (16%) and verbal knowledge (20%). This suggests that conflict monitoring mechanisms may play a more specialized role in reading comprehension, intervening relatively infrequently to resolve processing errors that would otherwise disrupt comprehension. This may explain why, in some fMRI studies of ‘naturalistic’ language comprehension, brain regions associated with conflict monitoring are infrequently reported (Fedorenko, 2014), particularly when the hemodynamic response is not time-locked to the onset of comprehension problems.

### Predictions and open questions

This domain-general conflict monitoring account raises some important questions and generates a number of testable predictions. First, do these results generalize to other types of linguistic error processing? In the current experiment, we examined individual differences in *semantic* error processing using local verb-argument mismatches. In future studies it will be important to determine whether domain-general conflict monitoring also influences sensitivity to other types of linguistic errors that are known to generate P600 effects (e.g. Osterhout & Holcomb, 1992; Vissers, Chwilla & Kolk, 2006). For example, readers with poor domain-general monitoring may be more likely to overlook misspellings or grammatical errors as they read, and they may experience difficulties when resolving lexical or syntactic ambiguities (Engelhardt, Nigg & Ferreira, 2017; Vuong & Martin, 2013). Similarly, future studies should also examine whether different types of linguistic anomalies also show a u-shaped relationship between conflict monitoring and the magnitude of the P600 response.

It will also be important to determine whether similar domain-general performance monitoring mechanisms are involved in the detection and resolution of more *global* discourse comprehension errors. During discourse comprehension, readers must encode abstract information that unfolds across multiple sentences, including thematic content, causal relationships, and character motivations (Baker & Anderson, 1982; Kinnunen & Vauras, 1995). Similar to local semantic anomalies, the detection of global conflicts can also indicate the onset of comprehension problems (*e.g. a vegetarian protagonist eating a hamburger*), which may prompt the reader to revise or reinterpret prior contextual information (Albrecht & O’Brien, 1993, Braasch, Rouet, Vibert & Britt, 2011; Hakala & O’Brien, 1995). Although related, we believe that the detection of global discourse anomalies may play an even more important role in successful text comprehension (Oakhill, Hartt & Samols, 2005), potentially relying on both domain-general conflict monitoring and working memory resources.

Given our current findings, it is also important to consider the relationship between error processing in *comprehension* and the error monitoring processes involved in language *production.* It is well established that speakers continually monitor their own production in order to prevent or correct unintended speech errors. Many aspects of error monitoring in language production are relatively automatic, involving the detection and resolution of internal representational conflicts within the production system (Nozari, Dell & Schwartz, 2011), which may be linked to certain domain-general executive functions (e.g. Altmann, Kempler, Andersen, 2001; Gollan, Smirnov, Salmon & Galasko, 2020). Here we suggest that, even more relevant to the comprehension error monitoring construct examined in the present study, are those aspects of error monitoring in language production that are less automatic, involving the conscious detection of errors and the initiation of downstream repairs (e.g. “*turn left* …. *uh, right*”). Traditional monitoring accounts of language production have argued that these latter types of error monitoring processes are carried out by the producer’s own comprehension system (monitoring through a perceptual loop, Levelt, 1983, see Gauvin & Hartsuiker, 2020 for a recent discussion). We suggest that linguistic error detection during *both* language comprehension and production may rely on a common set of computational mechanisms that are also engaged in aspects of domain-general conflict monitoring (Yu, Dayan & Cohen, 2009). In future studies it will be important to test this hypothesis more directly by examining the neural and behavioral correlates of error monitoring during both language comprehension and production.

### Neurobiology of Conflict Monitoring

The error monitoring account discussed in this study also makes predictions regarding the relationship between linguistic error processing and neurobiological measures of conflict monitoring. In previous studies, conflict-driven attention shifts have been associated with the phasic release of norepinephrine (NE) in the locus coeruleus (Aston-Jones & Cohen, 2005), which is thought to serve as an ‘orienting signal’ to interrupt the default task state (Yu & Dayan, 2005; Dayan & Yu, 2006; Bouret & Sara, 2005). The phasic release of NE has also been linked with positive-going ERP responses like the P300 (Nieuwenhuis, Aston-Jones & Cohen, 2005; Vazey, Moorman & Aston-Jones, 2018), which is thought to be functionally related to the P600 response (Coulson, King & Kutas, 1998; Osterhout, Kim & Kuperberg, 2012; Sassenhagen, Schlesewsky & Bornkessel-Schlesewsky, 2014; Sassenhagen & Fiebach, 2019). Also of relevance is a domain-general ERP component known as the “error positivity” or *Pe* (Falkenstein, Hohnsbein, Hoormann, & Blanke, 1990), which has been associated with the conscious, “metacognitive” recognition of errors (Nieuwenhuis, Ridderinkhof, Blom, Band, & Kok, 2001) and compensatory processes, such as additional information seeking, in the service of optimizing performance (Desender, Murphy, Boldt, Verguts, & Yeung, 2019). It will therefore be important for future studies to investigate whether linguistic error detection is associated with physiological markers of phasic NE release, such as pupil dilation (Sara, 2009), and whether P600 amplitudes are correlated with other positive-going ERP responses elicited in classic error monitoring tasks.

In fMRI studies, conflict-inducing tasks such as the Stroop and AX-CPT have been shown to activate a fronto-parietal network of ‘cognitive control regions’, including the anterior cingulate and lateral prefrontal cortex (van Veen & Carter, 2002; Niendam, et al., 2012). If linguistic conflict detection relies on the same neurocognitive mechanisms, then a similar set of brain regions should be recruited when comprehenders detect linguistic errors. Indeed, in fMRI studies, semantic anomalies and non-linguistic conflicts (*incongruent Stroop trials*) have been shown to activate overlapping regions of left inferior frontal cortex (van de Meerendonk, Rueschemeyer & Kolk, 2013; Ye & Zhou, 2009). Similar co-activations within this region have also been observed for sentences with syntactic ambiguities (Hsu, Jaeggi & Novick, 2017), which readers may initially interpret as syntactic errors (see Novick, Trueswell & Thompson-Schill, 2010; Nozari & Thompson-Schill, 2017 for reviews).

We should note, however, that this does not imply that these frontal regions contribute directly to the posterior P600 effect observed at the scalp surface (see van de Meerendonk, Rueschemeyer & Kolk, 2013 for a discussion). For example, frontal activity may reflect the initial *detection* of conflict, which then triggers additional compensatory mechanisms that are reflected in the P600 itself. Consistent with this idea, in a recent multimodal neuroimaging study (Wang, et al., 2021), semantic anomalies produced robust activations within left inferior frontal cortex and anterior cingulate, from 300-500ms after word onset. Critically, this activity was followed by a late reactivation of fusiform cortex (from 600-800ms) that more closely matched the timing of the P600 response. This late fusiform response may have reflected an orthographic re-analysis of the bottom-up input following an error.

### Practical applications

Our findings also have potential applications for the diagnosis and treatment of reading comprehension impairments. Reading comprehension problems are one of the most frequently observed deficits in children with learning disabilities (Lyon, 1995; Vaughn, Levy, Coleman, & Bos, 2002). In addition to problems with vocabulary and phonological awareness, poor readers often show impairments in their ability to actively monitor their comprehension as they read. For example, poor readers are less accurate at detecting linguistic anomalies (Garner, 1980; Hacker, 1997; Oakhill & Cain, 2012; Rubman & Salatas Waters, 2000), and, when comprehension errors *are* detected, these readers are less likely to engage in useful compensatory strategies such as re-reading (Ehrlich, Remond, Tardieu, 1999; Hacker, Dunlosky, & Graesser, 1998). Given our findings in adults, it is possible that children with poor domain-general conflict monitoring may be predisposed to develop reading comprehension problems (Cutting, Materek, Cole, Levine & Mahone, 2009). If this is the case, then early screening for conflict monitoring deficits could be useful tool for identifying at-risk readers, who may benefit from additional instruction or targeted interventions (Gerseten, Fuchs, Williams & Baker, 2001; Suggate, 2016).

Finally, a number of neuropsychiatric disorders that affect language processing have been linked to deficits in domain-general conflict monitoring, including schizophrenia (Kuperberg, 2010; Boudewyn, Carter & Swaab, 2012; Kerns et al., 2005; Lesh, Niendam, Minzenberg & Carter, 2010) and autism spectrum disorder (Agam, Joseph, Barton & Manoach, 2010; Solomon, Ozonoff, Cummings & Carter, 2008; South, Larson, Krauskopf & Clawson, 2010). Consistent with our current findings, individuals with these neurodevelopmental disorders also show marked impairments in linguistic error detection and the magnitude of the P600 response (Koolen, Vissers, Egger & Verhoevena, 2013; Kuperberg, McGuire & David, 1998; Kuperberg, Sitnikova, Goff & Holcomb, 2006). An important goal for future research will be to determine whether similar abnormalities in linguistic error processing are observed across diagnostic boundaries (Garvey et al., 2010) and whether they can be explained by domain-general conflict monitoring deficits.

## Footnotes

1 Several internal and external factors can influence the probability of generating a P600 response (Kuperberg, 2007). These include the nature of the experiment task (Geyer, Holcomb, Kuperberg & Perlmutter, 2006; Payne, Stites & Federmeier, 2019), the length and constraint of the prior linguistic context (Kuperberg, Brothers & Wlotko; 2020; Brothers, Wlotko, Warnke & Kuperberg, 2020; Nieuwland & Van Berkum, 2005), and, in impoverished contexts, the presence/absence of semantic attraction (e.g. Kuperberg, 2007; Kim & Osterhout, 2005).

2 In the psycholinguistic literature, the resolution of linguistic errors has been discussed within multiple frameworks that are often specific to the type of error encountered. For example, the syntactic parsing literature has focused on the triggers for reanalysis in garden-path sentences (single-stage vs. two-stage accounts, see Traxler, 2014), as well as distinctions between the “revision” of syntactic ambiguities and the “repair” of syntactically ill-formed sentences (e.g. Gouvea, Phillips, Kazanina, Poeppel, 2010; Kaan & Swaab, 2013; Vuong & Martin, 2014). The literature on semantic errors has also focused on different triggers for reanalysis (Kuperberg, 2007) and the potential resolution of anomalies through a noisy channel framework (Gibson, Bergen & Piantadosi, 2013). Here, we take a complementary and more general perspective, inspired by Kolk and colleagues, who proposed that these different forms of linguistic error processing are linked through a domain-general process of conflict monitoring (see Kolk & Chwilla, 2007; van de Meerendonk, Kolk, Chwilla & Vissers, 2009 for a discussion).

3 In control-demanding tasks, the presence of conflict can produce behavioral adjustments at multiple time scales, including the reactive re-allocation of attention within the current trial (Gehring, Goss, Coles, Meyer & Donchin, 1993), adaptations to optimize performance on subsequent trials (Gratton,Coles & Donchin, 1992, Botnovick, Nystrom, Fissell, Carter & Cohen, 1999), and longer-term learning that allows for the acquisition of new goals or task schemas (Botnivick, 2007).

4 Consistent with this suggestion, in our multiple regression analysis, we observed a marginally significant quadratic effect of verbal knowledge on semantic error detection (Verbal^2^: b = −.10, t = 1.79, *p* = 0.08). Specifically, in participants with very low verbal knowledge scores (the bottom tertile), error detection was less accurate (d’ = 2.3, SD = 0.5) than participants with higher verbal knowledge (d’ = 2.7, SD = 0.7).

## Notes

### Competing Interest Statement

The authors have declared no competing interest.

