## Supplementary Materials for "Domain-general conflict monitoring predicts neural and behavioral indices of linguistic error processing during reading comprehension"

***Task Order***

Participants completed all assessments in a single, two-hour session in the following order: Word Identification, Rapid automatized Naming, Rapid Automatized Switching, Woodcock Reading Mastery Test, TOWRE Sight Word Efficiency, North American Adult Reading Test, Subtract-Two Span, Listening Span, Kauffman Test of Educational Achievement, Manual Stroop, AX Continuous Performance Task, Operation Span, Reading Span, Author Recognition. Participants completed an additional set of tasks measuring verbal fluency, visual statistical learning, and phonemic awareness, which were outside the scope of the present study.

***Working Memory Capacity Tasks – additional information***

*Operation Span* - (Input Modality: *Visual*; Processing Task: *Math problem verification*; Memory task: *Letter Recall*; Span sizes: *3-7*; Trials per span*: 3*; Max score: *75*)

*Reading Span* - (Input Modality: *Visual*; Processing Task: *Sentence verification*; Memory task: *Letter Recall*; Span sizes: *3-7*; Trials per span*: 3*; Max score: *75*)

*Listening Span* - (Input Modality: *Auditory*; Processing Task: *Sentence verification*; Memory task: *Recall of sentence-final words*; Span sizes: *2-6*; Trials per span*: 3*; Max score: *60*)

*Subtract-Two Span* - (Input Modality: *Auditory*; Processing Task: *Subtract two from each digit*; Memory task: *Digit Recall*; Span sizes: *2-8*; Trials per span*: 5*; Max score: *175*)

***Conflict Monitoring Tasks – additional information***

<u>*AX-Continuous Performance Task*</u> **–**

Trial Numbers: *AX* = 105, *AY* = 15, *BX* = 15, *BY* = 15

Presentation Parameters: *Cue Duration* = 250ms, *Cue-Target Interval* = 500ms, *Target Duration* = 250ms, *Inter-trial Interval*: 500ms

Responses more than 750ms after target onset were classified as errors. We saw no significant differences in reaction time or accuracy when comparing BX and AY trials (*t*s < 1). This suggests participants has no difficulty maintaining cue information across a 500ms delay, and that this fast-paced version successfully minimized the role of context maintenance on performance.

<u>*Manual Stroop Task*</u> **–**

Trial Numbers: *Congruent* = 28, *Incongruent* = 28, *Neutral* = 28

Presentation Parameters: *Stimulus Duration* = until response, *Inter-trial Interval*: 500ms

Font colors: Black, Green, Red, Blue

The mapping between response keys and font colors was displayed at the top of the screen throughout the task. Participants received immediate visual feedback (*correct*/*incorrect*) after each response.

Table S1. Differences across the three versions of the ERP experiment

| | Trials per condition | Recording electrodes | Experimental + filler items | % High Constraint | # of Experimental Lists |
|---|---|---|---|---|---|
| Version A (N = 26) | 20 | 32 | 160 | 50% | 5 |
| Version B (N = 22) | 29 | 32 | 290 | 70% | 3 |
| Version C (N = 29) | 25 | 64 | 175 | 57% | 4 |

Note. We have previous reported data from larger samples of participants from Version A in Kuperberg, Brothers & Wlotko, 2020, Version B in Brothers, Wlotko, Warnke & Kuperberg, 2020; and Version C in Wang, et al., (in prep). Individuals who did not participate in additional neuropsychological testing were not included in the current analyses.

***Replication Experiment (N=37)***

In addition to our primary individual differences analysis (N=77), we also analyzed data from a separate ERP experiment (N=37) in which participants judged the plausibility of individual sentences that were either plausible or semantically anomalous (42 trials per condition). Participants in this sample did not complete the full individual differences battery, but they did complete a version of the AX Continuous Performance Task in the same experimental session. This AX-CPT task had a slightly longer cue-target SOA (750ms vs. 1000ms) and a larger number of trials (150 vs. 400) but produced similar patterns of behavioral results (AY accuracy = 81%, AY trial RT cost = 129ms).

In an attempt to replicate some of our primary findings from our main individual differences analysis, we combined our two AX-CPT behavioral measures into a single *conflict monitoring* score, and examined the relationship between these scores and 1) behavioral measures of linguistic error monitoring (plausibility d'), and 2) the amplitude of the P600 response (see Figure S1). Consistent with our prior behavioral findings we observed a robust, positive correlation between conflict monitoring and linguistic error detection abilities ($r(35) = 0.64$, $p < .001$). We also observed a similar inverted u-shaped relationship between conflict monitoring abilities and the amplitude of the P600 effect, although this effect was only marginally significant ($r(35) = 0.31$, $p = 0.07$) – perhaps due to the reduced sample size.

Table S2. Conflict monitoring effect sizes across experiments (*small* = 0.1; *medium* = 0.3; *large* = 0.5)

| | Correlation of interest | Effect Size (*r*) | 95% Interval [lower - upper] |
|---|---|---|---|
| Main Experiment (N=77) | Conflict Monitoring - Plausibility (d') | 0.29 | [0.07 - 0.48] |
| | Conflict Monitoring - P600 effect* | 0.40 | [0.19 - 0.57] |
| | Conflict Monitoring - Reading Comp. | 0.23 | [0.00 - 0.43] |
| Replication (N=37) | Conflict Monitoring - Plausibility (d') | 0.64 | [0.40 - 0.80] |
| | Conflict Monitoring - P600 effect* | 0.31 | [-0.01 – 0.58] |

* Quadratic effect; Reading Comp. = Reading Comprehension Tasks

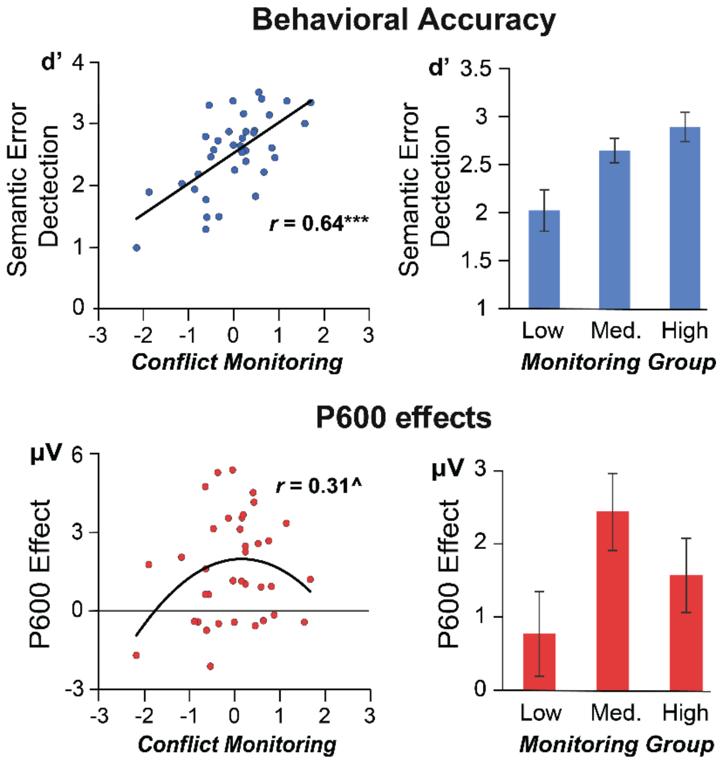


Figure 1. Relationship between conflict monitoring scores and behavioral and neural measures of semantic error processing in an independent sample. Error bars represent ±1 SEM. *** $p < .0001$, ^ $p = .07$

*Replication Experiment – Additional Information*

Participants were 37 adults (19 female; average age = 24) recruited from Tufts University and the surrounding community. Inclusion criteria and consent information were identical to previous ERP experiment. Participants read single sentences presented one word at a time in the center of a computer screen. Half of the sentences were plausible, and half were semantically anomalous, with anomalous items generated by swapping critical words across sentences (*Joan fed the baby some warm peas/offices from the supermarket)*. At the end of each trial, participants responded via button-press whether the preceding sentence was plausible or anomalous. EEG was recorded simultaneously from 32 electrode sites. Acquisition and data analysis methods were identical to those in the previous experiments, and P600 effects were analyzed in the same central-posterior region-of-interest from 600 to 1000ms after critical word onset (anomalous vs. plausible: 1.6µV, $t(36) = 4.95$, $p < .0001$).
